# pylimma: a faithful, AnnData-native Python port of R limma for differential expression analysis

**DOI:** 10.64898/2026.07.06.736732

**Authors:** JF Mulvey

## Abstract

*pylimma* is a faithful Python port of *limma*, intended to bring one of the most widely used tools for differential expression analysis to the developing Python ecosystem for transcriptomics and proteomics. We validated *pylimma* against the existing R implementation through 227 function-level comparisons and across six real world datasets spanning microarray, RNAseq, proteomics and single-cell transcriptomics. *pylimma* reproduces *limma*’s numerical output to a median agreement of 13 significant figures and calls identical sets of differentially expressed features and gene sets. This supports its use as a drop-in replacement for the R implementation.

## Introduction

The *limma* package for R has become a foundational tool for analysis of -omics datasets. Whilst it was originally developed for the analysis of microarray data, it now receives widespread use across microarray, RNAseq and proteomics data types. As a crude metric for the use that this tool has received, the paper introducing the application of *limma* to bulk RNAseq data^1^ has alone been cited 43,144 times within the OpenAlex dataset^2^ at the time of writing, making it the 27th most cited paper within the field of molecular biology and a *bona fide* citation classic. The current version represents the culmination of over 20 years of development from Gordon Smyth and colleagues, who originally proposed the idea of empirical Bayes shrinkage of gene-wise variances in 2004^3^. Key subsequent developments include (but are not limited to) the application of quality weights across samples^4^, fold-change aware hypothesis testing^5^, gene set testing accounting for correlation between genes^6^, and the ability to test for differential transcript usage^7^. The current version contains ∼15,000 lines of R code plus a small amount of C, and remains maintained on Bioconductor by Gordon Smyth’s team at WEHI.

Whilst once the applied -omics community was once coalesced around the R ecosystem, it is increasingly split between R and python. scRNAseq tooling for example has increasingly crystallised around Python, driven by initiatives such as the scverse project^8^, the AnnData data structure^9^ and multimodal extensions such as MuData that allows the flexible use of AnnData structures for a range of multimodal data types^10^. Practitioners who want to employ *limma*’s empirical-Bayes moderation - which is consistently a top-tier method in benchmarks of pseudobulk differential expression analysis methods^11,12^ - must currently choose between switching languages mid-pipeline or accepting an alternative method whose statistical properities may not be as desirable. Beyond single cell transcriptomics, the field of proteomics is also increasingly adopting python based workflows especially as technological advances mean that measuring (many) single cells or spatial areas by mass spectrometry has become feasible even beyond specialist laboratories. The ProteoPy^13^ and openDVP^14^ packages are beginning initiatives to help codify this, although the lack of options for statistical testing is a major drawback to those currently considering migrating from the existing R ecosystem.

There is therefore a clear need for access to the statistical methods made available within the R *limma* package for the python community. A common solution has has been to use rpy2, which allows R processes to be called from within python. This requires an external R dependency. Alternatively inMoose ports a small subset of 17 *limma* functions to python for native use^15^. A small number of other projects exist to port *limma* functionality to python, but none cover more than the basic pipeline or present demonstrations of their parity to the output of the R *limma* package. Moreover, none of these are designed to seamless utilise the AnnData structure that has become *the* central component of the current python ecosystem.

Here, we present and validate *pylimma*, a faithful and comprehensive end-to-end python port of the R *limma* package. This preserves the user interface of the l*imma* package such that current users of *limma* are able to seamlessly migrate, and as we demonstrate produces strict parity in output against the R *limma*. It extends the functionality of *limma* to allow full native compatibility with AnnData objects. The package is thoroughly documented, and includes 6 example applications on real datasets covering microarrays, bulk RNAseq, assessment of differential transcript usage, mass spectrometry proteomics and pseudobulk analysis of single cell RNAseq.

## Results

*pylimma* contains 17,717 lines of code, which maintain the same user interface as the *limma* package with the exception of the use of snake case rather than camel case (Figure 1A). Parameter names and defaults track those in limma 3.66.0. *pylimma* furthermore allows native input from, computations upon and output to AnnData objects, whilst also porting the familiar EList data structure and compatibility with flat matrices such as NumPy arrays and pandas DataFrames (Figure 1B). *pylimma* covers linear modelling with empirical Bayes moderation, voom precision weights, normalisation between arrays, background correction, removeBatchEffect, duplicate correlation, array weights, TREAT, gene-set testing (camera/roast/mroast/fry/romer/geneSetTest), differential splicing, diagnostic plots and the supporting numerical utilities. We do not port two-colour-array containers (`RGList, MAList`), `ExpressionSet`, illuminaio readers and the affy-driven probe-summarisation pipeline (Fig 1C, full details in Table S1).

**Figure 1.**
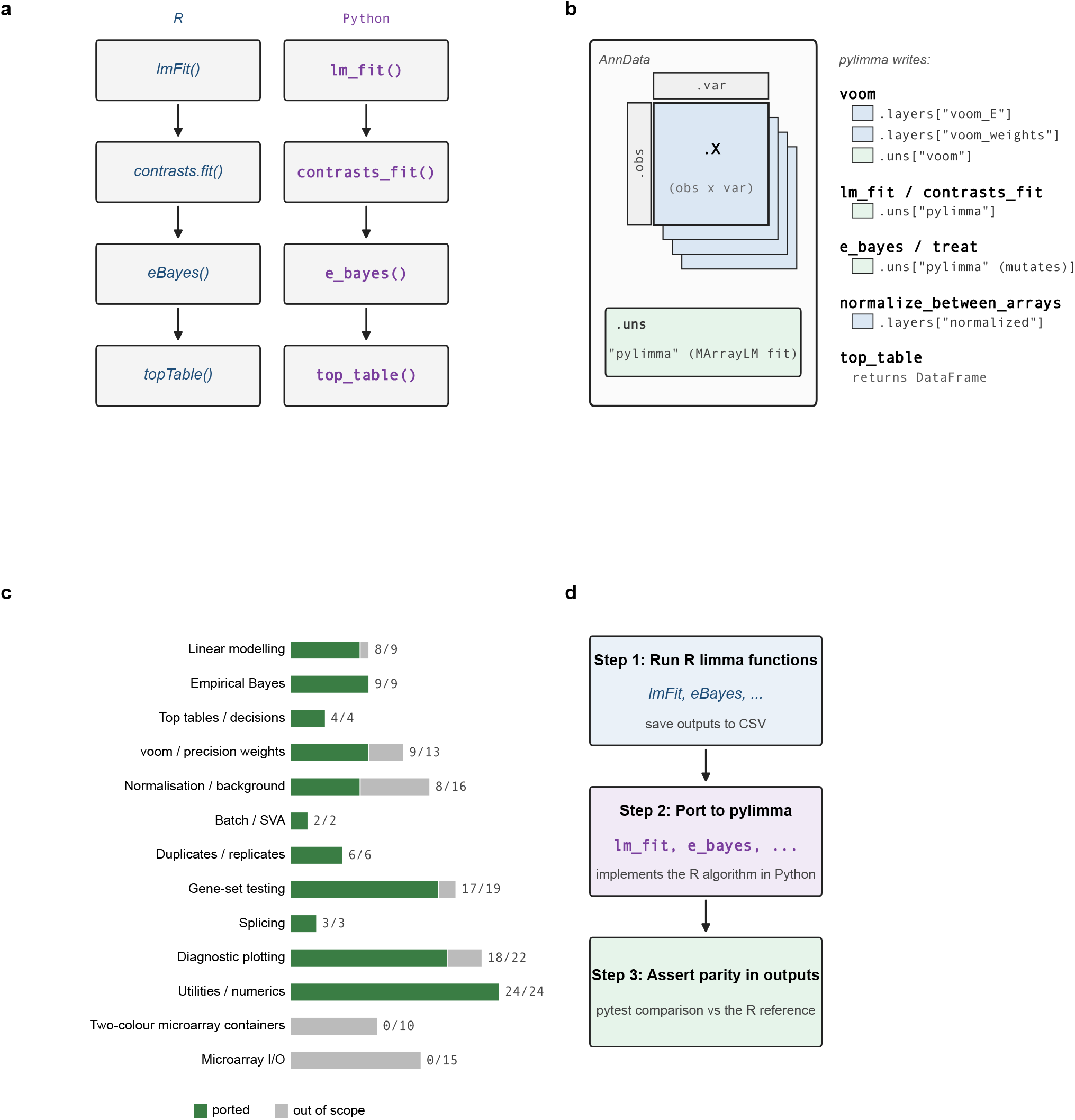
Scope, architecture and validation strategy of *pylimma*. **A**. Pipeline diagram of the canonical *limma* workflow and the equivalent in *pylimma*. User interface is consistent across both tools, beyond the use of camel case in *limma* and snake case in *pylimma*. **B**. Polymorphic input/output dispatch allows *pylimma* to use not only conventional EList data structure, but also natively compute on and interact with AnnData structures. The schematic shows where the output of commonly utilised functions are stored within the AnnData container. **C**. Summary of the functional coverage of *limma* contained within *pylimma*. Microarray specific data readers and two-colour array specific functionality were determined to be out of scope for the initial version. **D**. Schematic of the implementation loop. An R script using *limma* 3.66.0 was used to generate a csv output covering a diverse set of use cases for each function. The corresponding *pylimma* function was then written, and iterated upon until parity in output with the R fixture was obtained.

Community contributions to extend the port to cover this functionality are welcome. The port was performed by first designing small input datasets that could be used to exercise the distinct code paths and input options within each of *limma*’s functions in turn. *limma* was run against these, the function ported to python and the new python function’s output compared against the *limma* output. The python function was iterated upon until an acceptable degree of parity was met (Figure 1D).

To confirm that *pylimma* accurately reproduces *limma*’s computations, we compared the output of *pylimma* and *limma* element-by-element. Reference outputs for every ported function were generated in R using *limma* 3.66.0 on simulated inputs chosen to exercise each function’s code paths. *pylimma* was run with identical inputs. For each of 227 comparisons we recorded the worst relative difference across all elements of the output (Figure 2A). The median of these values was 10^-13.7^, equivalent to agreement to ∼13 significant figures, whilst no comparison exceeded 2.4×10^-4^. That largest difference arose in normalizeVSN, whose profile likelihood is near-flat along one direction so different optimisers settle at marginally different points. Here the outputs are nonetheless visually identical (R^2^ = 1.000000; Figure 2A inset, with data point in main panel highlighted in red).

**Figure 2.**
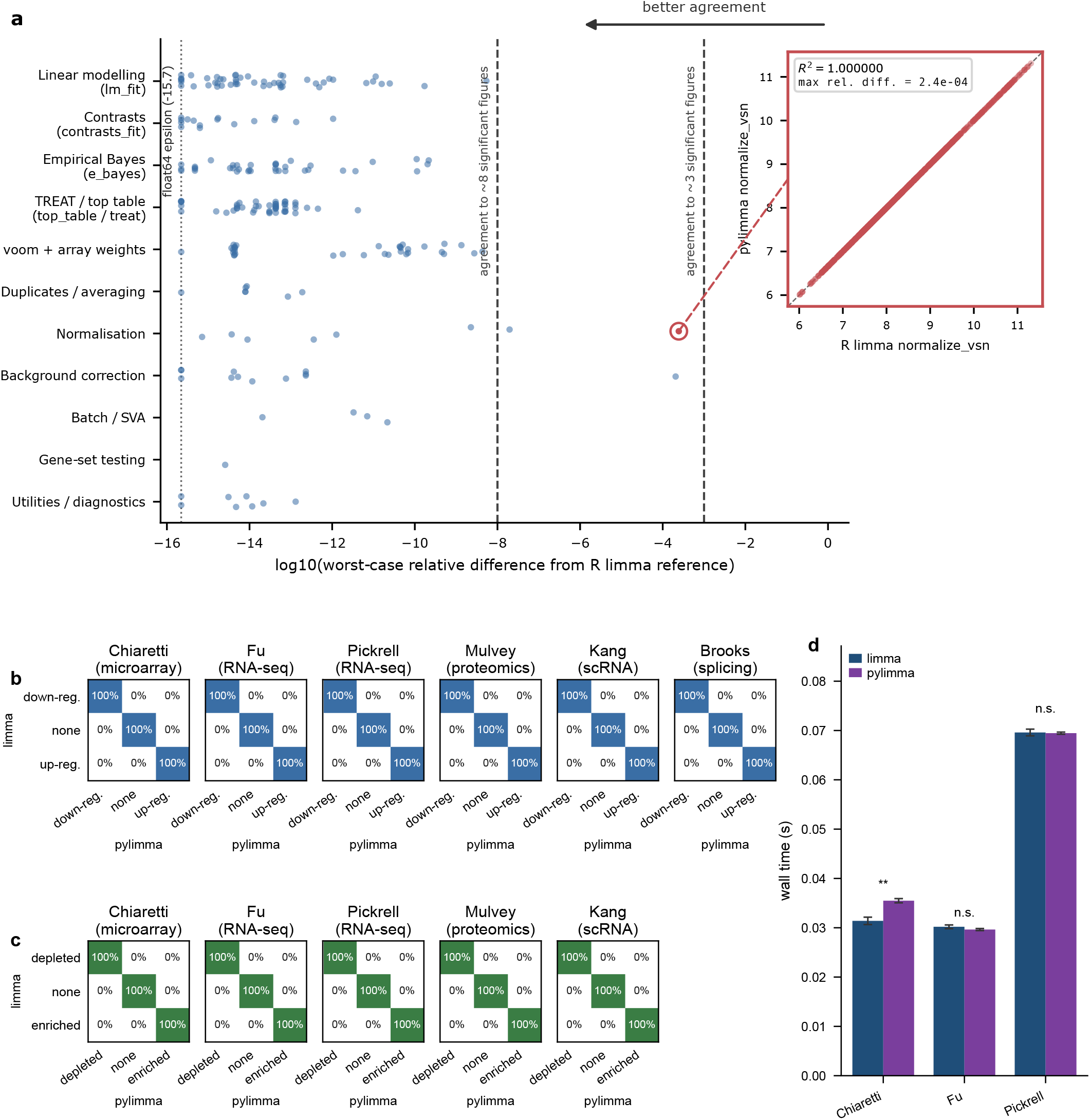
*pylimma* demonstrates numerical parity in outputs with R *limma*. **A**. Comparison of numerical output between *pylimma* and *limma* 3.66.0. Each datapoint represents a comparison between a different set of input parameters that are chosen to exercise the distinct code paths and options of each function. We then compare each numeric output and show the maximum relative error between *pylimma* and *limma*. The x-axis is presented on a log10 scale, with a vertical dotted line at -15.7 indicating 64 bit floating point error. The inset shows the correlation between R *limma* and *pylimma* for the test which has the greatest tolerance between *limma* and *pylimma*, which nonetheless shows an R^2^ of 1.000000 (correct to 6 decimal places). **B**. 6 real datasets were selected to assess practically relevant output of *pylimma* across a diverse range of use cases. For each dataset a confusion matrix is shown showing the percentage of genes/proteins/transcripts as appropriate that were called a significantly up- or down-regulated between conditions in *limma* and *pylimma*. **C**. A gene set enrichment analysis was performed upon protein coding genes or proeins using the camera method. A confusion matrix shows the agreement in gene sets called as significantly up- or down-regulated between conditions when running the entire pipeline in either *limma* or *pylimma* **D**. Computational time of the core lm_fit → e_bayes pipeline within each package (*n* = 5; Welch t-test, ** *p* < 0.01, n.s. not significant) for the three bulk datasets.

Secondly, we provide a practical comparison of real world performance utilising pre-existing datasets covering a range of different experimental methods (microarray^16^, RNAseq^17,18^, single cell RNAseq^19^, mass spectrometry proteomics^20^), making use of conventional^16,19,20^, voom^17,18^ and differential splicing^21^ pipelines. Calling of significantly up- or down-regulated genes was identical between *pylimma* and *limma* (Figure 2B), as was the case when running the full pipeline to analyse differentially up- or down-regulated gene sets with camera (Figure 2C). No discordant calls were observed across any of the features tested across the six datasets. This was achieved with no difference or a very minor increase in computational time upon the core lm_fit → e_bayes pipeline. The magnitude of this any difference is not practically significant, with all computations from either package complete within 72 ms.

## Discussion

*pylimma* is a comprehensive and validated Python port of R *limma*. Beyond faithfully porting existing functionality, we also allow AnnData-native use in order to facilitate integration within the existing scverse python ecosystem. Consistent with the nature of *pylimma* as a derivative work of *limma*, we encourage users of the tools to cite the original statistical papers for the relevant methods to ensure that the original authors receive credit for their ideas in addition to the *pylimma* implementation. This follows the conventions established in previous ports of analytic software such as PyDESeq2^22^ and most recently edgePython^23^.

In this manuscript we provided a thorough validation of *pylimma*’s numerical output in order to demonstrate that *limma* and *pylimma* can be used in an interchangable manner. Beyond the analysis presented here the package contains the full analysis pipeline for all 6 datasets summarised here, in order to aid new users to *limma* to start directly within *pylimma*.

*pylimma* enables a diverse set of analyses to be performed directly within python. Most notable amongst the novel use cases that this is designed to help enable is the pseudobulk differential expression analysis of single cell RNAseq data. Similarly, analysis of mass spectrometry based proteomics can be performed using the AnnData structure directly with an example present from Mulvey et al.^20^. We note that the contemporary edgePython implementation of edgeR allows only exporting of DGEList results via a to_anndata export step rather than allowing the differential expression step to live *inside* an AnnData object as *pylimma* enables.

Every effort has been made to ensure parity with R *limma* in output, interface and scope. By releasing the *pylimma* source code openly we hope to let the community identify any behaviour that falls outside our validation and testing strategy: that the many eyes of open development inevitably surface issues that no single test suite can anticipate^24^. We welcome such scrutiny in use, and the contributions that follow from it. In this vein, porting *limma* line-by-line itself involved close scrutiny of the *limma* codebase and in doing so we identified two minor bugs in *limma*. In genas, a coefficient re-centring that *limma* clearly intends is silently discarded by a partial-matching quirk of R’s $<-operator. In voom, user-supplied design and weight slots are dropped during EList coercion. Where they arise *pylimma* reproduces the intended rather than the literal behaviour. That only two such edge cases surfaced across a package of *limma*’s age, size and scope is a testament to its quality. The remaining differences that we are aware of are not errors in either implementation but consequences of the two languages’ numerical libraries. For example, the rotation-based gene-set tests (within roast, mroast and romer) reproduce every deterministic quantity exactly. However they draw their rotations from a different generator (NumPy’s PCG64 vs R’s Mersenne Twister) and these cannot be aligned from a shared seed. Their Monte-Carlo *p*-values therefore agree only to within sampling error. We regard these as functionally equivalent to limma. We note that the tolerance by which we match the ouput of *limma* is often much more precise than the drift of behaviour between different versions of *limma* itself. For example, the voom pipelines disagree between 3.66.0 and 3.54.2 at percent-level (up to ∼3% in our hands) because *limma*’s own voom/LOWESS code changed between these versions. By way of another comparison point, the contemporary *edgePython* port^23^ validates against *edgeR* at a relative tolerance of 1e-3, putting *pylimma* >4x better tolerance for our worst case scenario with the reference R implementation than *edgePython*.

Lastly, we note that substantial proportions of key functions in R *limma* have been written in C in order to improve computational performance. To date no effort has been made to make *pylimma* more performant and it is thus currently slightly slower than R limma on several pipelines beyond the core `lm_fit()` and `e_bayes()` presented here. The voom mean-variance LOWESS (R has hand-written C; pylimma calls `statsmodels.nonparametric.lowess)` is the obvious target for future performance focused updates.

To conclude, *pylimma* provides the python community with a validated, interface-faithful port of R *limma*’s empirical Bayes framework, natively integrated with the AnnData structures that underpin the scverse ecosystem. We have aimed to be as transparent about the port’s limitations as we have been rigorous in validating its parity such that new users are able to trust its output enough to use it upon their own datasets. We hope that *pylimma* removes a long standing barrier for those who want access to *limma*’s statistical machinery without leaving python.

## Methods

### Source and licensing

*pylimma* was ported from R limma 3.66.0 and is such released under GPL-3.0-or-later in conformance with limma’s GPL (>=2) licence. R contributors to the original source files have been acknowledged in headers of the relevant *pylimma* files throughout the package. Copyright licencing from upstream dependencies of *limma* from which content has also been ported as part of the *pylimma* pakcage (MASS, statmod, base R splines, affy) are also recorded within file headers. Their work is also acknowledged within the ‘NOTICE’ distributed within *pylimma*. All analysis in this manuscript was performed using *pylimma* v0.1.0.

### Implementation

The line-level Python was written primarily by Claude (Opus 4.5-4.7, Anthropic, USA) under the direction and review of the author, who carries full responsibility for the port.

### Validation against *limma* output

Numerical parity with R limma (v3.66.0; R 4.5.2) was assessed on simulated data. A single R script generated 227 reference outputs from small synthetic inputs spanning the regimes each function required. Examples include two- and three-group linear-model designs, a matrix with missing values, an unequal-residual-df case, a blocked duplicate-correlation design and negative-binomial counts for voom. Each pylimma function was run on the identical input and its output compared element-wise to the R reference. For every assertion we recorded the worst-case relative difference, max_i_ |xi^py^ − x_i_^R^| / |x_i_^R^|. No hard threshold was set for the parity in precision, since some calculations within package functions are closed form whilst others rely on iterative routines that accumulate per-step rounding error (LOWESS smoothing inside voom, Fisher scoring inside `array_weights` and `duplicate_correlation`, normexp parameter optimisation).

### Computational Performance

Computational performance was assessed by running the standard lm_fit() and e_bayes() pipeline 5 times, and comparing mean differences between *limma* and *pylimma*.

### Statistics

Data were analysed using python 3.11, and are shown using the mean ± SEM.

## Acknowledgements

We would like to acknowledge the work of Gordon Smyth and the limma team over more than two decades in developing and maintaining the R limma package. All statistical credit for the methods behind pylimma belongs to these authors, including but not limited to Yifang Hu, Matthew Ritchie, Jeremy Silver, James Wettenhall, Davis McCarthy, Di Wu, Wei Shi, Belinda Phipson, Aaron Lun, Natalie Thorne, Alicia Oshlack, Carolyn de Graaf, Yunshun Chen, Goknur Giner, Mette Langaas, Egil Ferkingstad, Marcus Davy, Francois Pepin, Dongseok Choi, Charity Law, Mengbo Li and Lizhong Chen. We also acknowledge the authors of R packages upstream of limma whose algorithms pylimma also ports, most notably MASS (Brian Ripley, Bill Venables) for `rlm`; statmod (Gordon Smyth, Lizhong Chen) for `mixedModel2Fit` and `glmgam.fit;` base R splines (R Core Team, original code by Douglas Bates and William Venables) for ns(); affy (Rafael Irizarry, Laurent Gautier, Benjamin Bolstad and contributors) for `bg.parameters`.

## Data and Code Availability

pylimma is open source software released under the GPLv3 license. pylimma is available at github.com/john-mulvey/pylimma and can be installed via PyPI: `pip install pylimma`.

Datasets used to demonstrate pylimma in this manuscript are derived from previously published studies and are redistributed within pylimma where possible for the convenience of users.

